# AliMarko: A Novel Tool for Eukaryotic Virus Identification Using Expert-Guided Approach

**DOI:** 10.1101/2024.07.19.603887

**Authors:** Nikolay Popov, Anastasia Evdokimova, Ignat Sonets, Maria Molchanova, Vera Panova, Elena Korneenko, Alexander Manolov, Elena Ilina

## Abstract

Metagenomic sequencing is a valuable tool for studying viral diversity in biological samples. Analyzing this data is complex due to the high variability of viral genomes and their low representation in databases. We present the Alimarko pipeline, designed to streamline virus identification in metagenomic data. A key feature of our tool is the focus on the interpretability of findings: results are provided with tabular and visual information to help determine the confidence level in the identified viral sequences.

The pipeline employs two approaches for identifying viral sequences: mapping to reference genomes and de novo assembly followed by the application of Hidden Markov Models (HMM). Additionally, it includes a step for phylogenetic analysis, which constructs a phylogenetic tree to determine the evolutionary relationships with reference sequences. We also emphasize reducing false-positive results. Reads related to cellular organisms are computationally depleted, and the identified viral sequences are checked against a list of potential contaminants. The output is an HTML document containing visualizations and tabular information designed to assist researchers in making informed decisions about the presence of viruses. Using our pipeline for total RNA sequencing of bat feces, we identified a range of viruses and rapidly determined the validity and phylogenetic relationships of the findings to known sequences with the aid of reports generated by AliMarko.

## Introduction

Viral infections, especially of a zoonotic nature, are an important source of human infections [1]. Viruses have historically been responsible for numerous epidemics. The risk of viral zoonosis outbreaks is likely to increase due to climate change, which can facilitate the emergence and spread of these diseases . Effective monitoring of viral populations is crucial for preventing and responding to epidemics, and this requires the development of robust methods for identifying and characterizing viruses [2].

Viruses are inherently complex and challenging objects of study due to their genetic heterogeneity and the absence of a single, universal single copy genes that could serve as a marker for their identification and classification (similar to the 16S ribosomal RNA gene in bacteria) [3]. Instead they feature various hallmark genes that permit the identification of particular known viral families [4, 5]. The amplification of hallmark genes poses a significant challenge, primarily due to their high sequence divergence and large number. This limitation necessitates the use of metagenomic approaches for analyzing viral communities, which enable researchers to overcome the constraints of gene amplification [6].

The analysis of viral communities using metagenomic approaches is further complicated by the fact that viral sequences may be present at low abundance, making them more susceptible to contamination and increasing the risk of false-positive results. Contaminants may distort results of a research and cause wrong conclusions [7]. For example, contaminating ATCV-1 [8] and XMRV [9] sequences in human samples caused misassociation of them with alterations in cognitive function and chronic fatigue syndrome respectively [10, 11]. Asplund et al revealed an association between a wide range of viral sequences and reagents used in library preparation [12]. Other works report contaminations coming from RNA purification kits [13].

Several utilities are available for detecting viral sequences in metagenomes [14]. A commonly employed approach involves local sequence alignment against a reference database using the BLAST tool to identify viral sequences, or utilizing BLAST-based tools such as VirusSeeker [15]. Another widely used method is the mapping of short sequences to viral genomes, with BWA [16] and Bowtie2 [17] being two popular tools employed in this context. Additionally, Hidden Markov Models (HMMs) are used to recognize viral protein sequences, with HMMer3 being the most widely used instrument for this approach. A similar method involves the use of Position-Specific Scoring Matrices (PSSMs), which enable efficient computational searches for ungapped matches [18]. An alternative approach to detect viral sequences involves analyzing the k-mer (short sub-sequences of fixed length) spectrum. Kraken2 is a prominent tool that implements this approach [19].

Some tools use a combination of methods to leverage the advantages of different approaches. VirSorter2 uses a combination of HMM and common features analysis (like GC content and gene density) [20]. VirFinder uses a combination of k-mers and SVM and returns likelihood of being a viral sequence [21].

In recent years, several tools have been developed for identifying virus sequences using recurrent or convolutional neural networks, such as ViBE [22] and DeepVirFInder [23]. These models can be computationally intensive, which may limit their applicability.

To summarize, a number of tools have already been developed for the automatic identification of viral sequences. However, the complexity of these algorithms and the ambiguity in setting appropriate thresholds can pose significant challenges for practicing biologists and epidemiologists. Researchers often find themselves needing to verify the findings of automated tools to ensure the reliability of the results. Our approach focuses on providing the solution that helps researchers confidently assess and validate the outputs of automated systems and to filter false-positive findings.

Bats are known as a significant reservoir of a wide range of zoonotic viruses, including those from the Lyssaviridae, Filoviridae, Paramyxoviridae, and other families, which can cause severe diseases in humans [24]. Their unique characteristics, such as long lifespan, social behavior, and wide geographic distribution, make them a natural hub for this diverse range of viruses. Metagenomic sequencing has enabled researchers to explore the viral diversity of bats, revealing a vast number of unknown viruses [25]. The study of bat viruses is crucial for early detection and monitoring of potential health threats.

Metagenomic sequencing datasets from bat samples contain a large number of viral sequences, the analysis of which requires specialized tools that can accelerate data analysis and evaluate the reliability of results for datasets with a broad repertoire of viruses.

## Results

### Pipeline overview

The pipeline takes raw read files as input, performs quality filtering, and depletes reads from cellular organisms. Following these preprocessing steps, two main types of analyses are conducted: (1) mapping to reference genomes and (2) de novo assembly of contigs followed by HMM analysis and phylogenetic analysis. The pipeline generates an HTML report that provides users with comprehensive information, including statistics on the results of each module, presented in the form of tables and illustrations (Figure 1).

**Figure 1.**
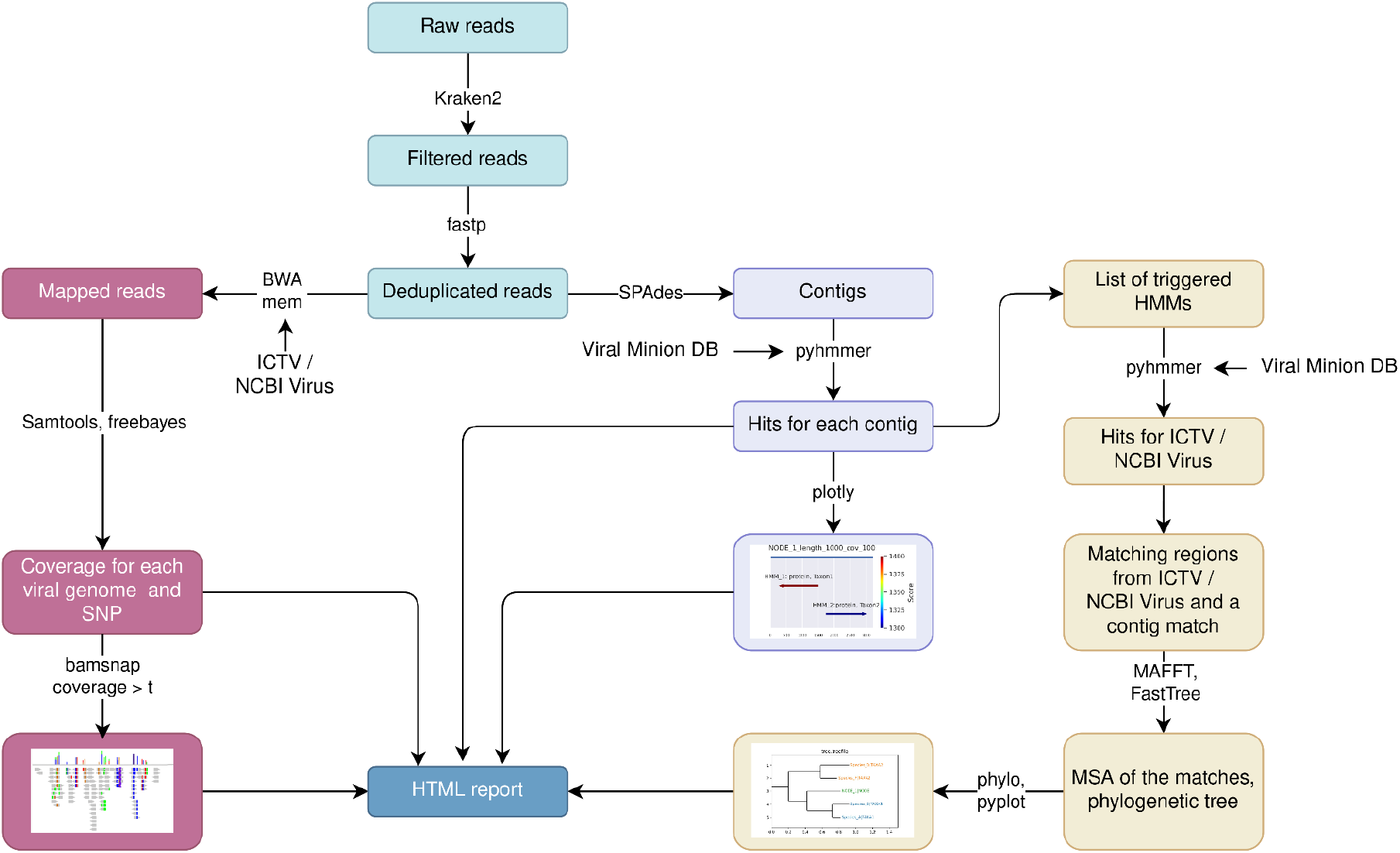
The schematic representation of the AliMarko pipeline. Two main types of analysis are performed: read mapping to reference genomes and HMM analysis of assembled contigs followed by phylogenetic analysis.

Depletion of reads from cellular organisms is achieved using Kraken2 with a custom database (see Methods). Ignoring this step can result in numerous false positives results, particularly due to nonspecific read mappings.

For mapping, we use reference genomes included in the Virus Metadata Resource from the International Committee on Taxonomy of Viruses (ICTV) [26]. ICTV develops a universal taxonomic scheme of viruses and provides access to VMR, which includes current taxonomy, information about host organisms, and representative genomes. This metadata is included in the report.

Visualization of read mapping to reference genomes helps assess the coverage width, the uniformity of coverage depth, and the density of single nucleotide substitutions. It is especially useful if the virus in the sample is similar to those present in the genomic database.

After the mapping, the correspondence of references to certain taxonomic groups is established according to the provided table (VMR) by custom script. The script takes into account that viral genomes can have several fragments, which act as different references when mapping. The script calculates average mapping characteristics of a virus taking into account length of its genome fragments.

We also provide information on the potential contaminant nature of a finding. For this, we use the information provided by Asplund et al. [12]. If a genome in the mapping results exhibits high similarity to a laboratory-component-associated sequence, it is flagged with red color and accompanied by a tooltip providing information about potential contaminant nature of the sequence.

The HMM module identifies conservative amino acid patterns specific to various viral groups. This component detects the viral origin of contigs and aids experts in result interpretation by visualizing HMM matches to the contig and displaying a phylogenetic tree of amino acid sequences matched with others in the selected reference database. To reduce false-positive results, we established a threshold for each HMM to enhance the accuracy of our detections.

The phylogenetic tree’s structure may confirm the viral nature of the match and identify the phylogenetically closest reference in the database. A consistent tree structure that aligns with accepted taxonomy, along with the contig fitting well within a specific clade, enables us to make a reliable conclusion. Conversely, weak phylogenetic signals, conflicting tree topology with taxonomy, or long branch length may suggest a false-positive result or a finding of a new viral representative. Due to the trees are made by a fragment of the genome, the topology of a tree may debate from the accepted taxonomy.

The pipeline supports batch analysis of multiple files, generating a comprehensive HTML report for an overview of results. Experts can then delve into specific sample reports for detailed insights.

### AliMarko reports

#### One sample report

After processing a file, AliMarko returns a sample’s HTML report. The sample’s HTML report includes tabular and graphical representations of analysis results (Figure 2, HTML S1)

**Figure 2.**
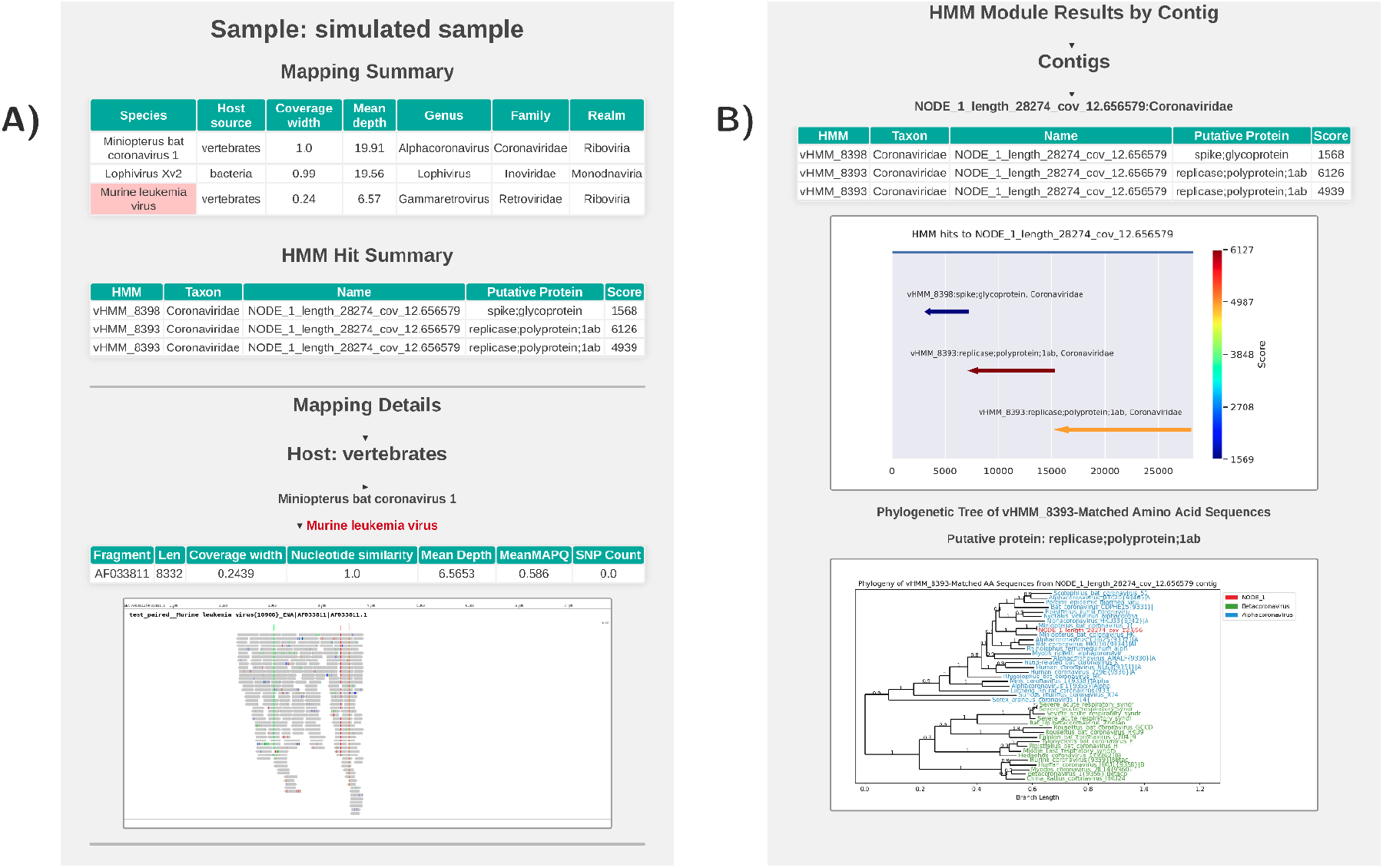
Screenshot of the AliMarko report for a sample. In the HTML report, the content of Figure B is located below the content of Figure A. Figure A: The report contains, from bottom to top, a comprehensive table with the results of the mapping module (Mapping General Results), followed by the results of the HMM module (General HMM Results). These are succeeded by the mapping details, which comprise several tabs for different assigned hosts. Upon opening the virus tab, a table displaying the mapping parameters and visualization is presented. If there is information on potential contamination for the reference genome, its name is highlighted in red. Figure B: Details for the HMM module. Each contig has its own tab, providing information on HMM hits on the contig, accompanied by a visualization of the hits. For each hit, a phylogenetic tree is constructed based on amino acid sequences. On the tree, reference sequences are colored according to taxonomic information.

At the top of the report, a table is displayed, which summarizes the results of the mapping module (Figure 2A). This table provides a comprehensive view of the mapping statistics with each row corresponding to a specific viral species. The table includes essential information such as the width and mean depth of coverage on the viral genome, host information and the taxonomic classification of the virus . The viruses are arranged in descending order of coverage width, thereby enabling a facile identification of the viral species with the most extensive coverage.

Below the mapping module table, a second table is presented, which summarizes the results of the HMM module (Figure 2A). Each row in this table corresponds to a single HMM hit on an assembled contig. The table includes essential information such as the HMM name, its meta-information, the contig name, as well as the score assigned to the hit and its relation to the threshold score (normalized score). The hits are arranged in descending order of normalized score, allowing for a straightforward identification of the most significant hits.

Following the tabular results, the report presents visualizations of the mapping data (Figure 2B). These visualizations are organized by the predicted host, as specified in the meta-information (at the kingdom level in VMR). Each virus has a separate tab, which, when expanded, reveals a table providing detailed information on the mapping parameters for each segment of the virus. The table includes essential information such as the width and depth of coverage, as well as the number of identified SNPs. Below the table, a visualization of the mapping data is presented, offering a graphical representation of the read distribution across the genome and the distribution of SNPs.

Below the mapping module details, the report presents the results of the HMM module in a tabular format (Figure 2B). Each contig is represented by a separate tab, which, when opened, reveals a comprehensive table summarizing the HMM hits against the contig. The table includes essential information such as the score obtained, meta-information about the HMM, and the normalized score. Additionally, a visualization of the HMM hits is provided for each contig, displaying the contig’s length in nucleotides and the corresponding HMM hits. Each hit is graphically represented by an arrow, whose boundaries and direction reflect the region and direction of translation that the HMM hit. The color of the arrow corresponds to the score assigned to the translation.

In the case presented in Figure 2B three HMMs from the same family matched the presented contig with a high score. The presence of hits on the polyprotein and the spike protein of *Coronaviridae* suggests a taxonomic affiliation of the contig and the functional potential of its sequence.

Below the HMM hit visualizations, the report presents phylogenetic tree visualizations for each HMM hit. These trees are constructed by combining the amino acid sequence of the hit with the amino acid sequences of translations from reference nucleotide sequences that also triggered this HMM. By default, the reference sequences are color-coded according to their genus classification, as determined by VMR. The phylogenetic trees provide a graphical representation of the relationships between the hit and the reference sequences, facilitating the evaluation of the HMM hit’s accuracy, its connections to known sequences, and the signal strength in the hit’s sequence.

#### Multisample report

If the pipeline receives several FASTQ files and processes them simultaneously.

In the multisample HTML report, an overview of the results is provided, giving a summary of the findings of alignment to the reference database and scanning the data by HMM. For a more detailed analysis of specific results, researchers can access the HTML report dedicated to a particular sample. (Figure 3A, B)

**Figure 3.**
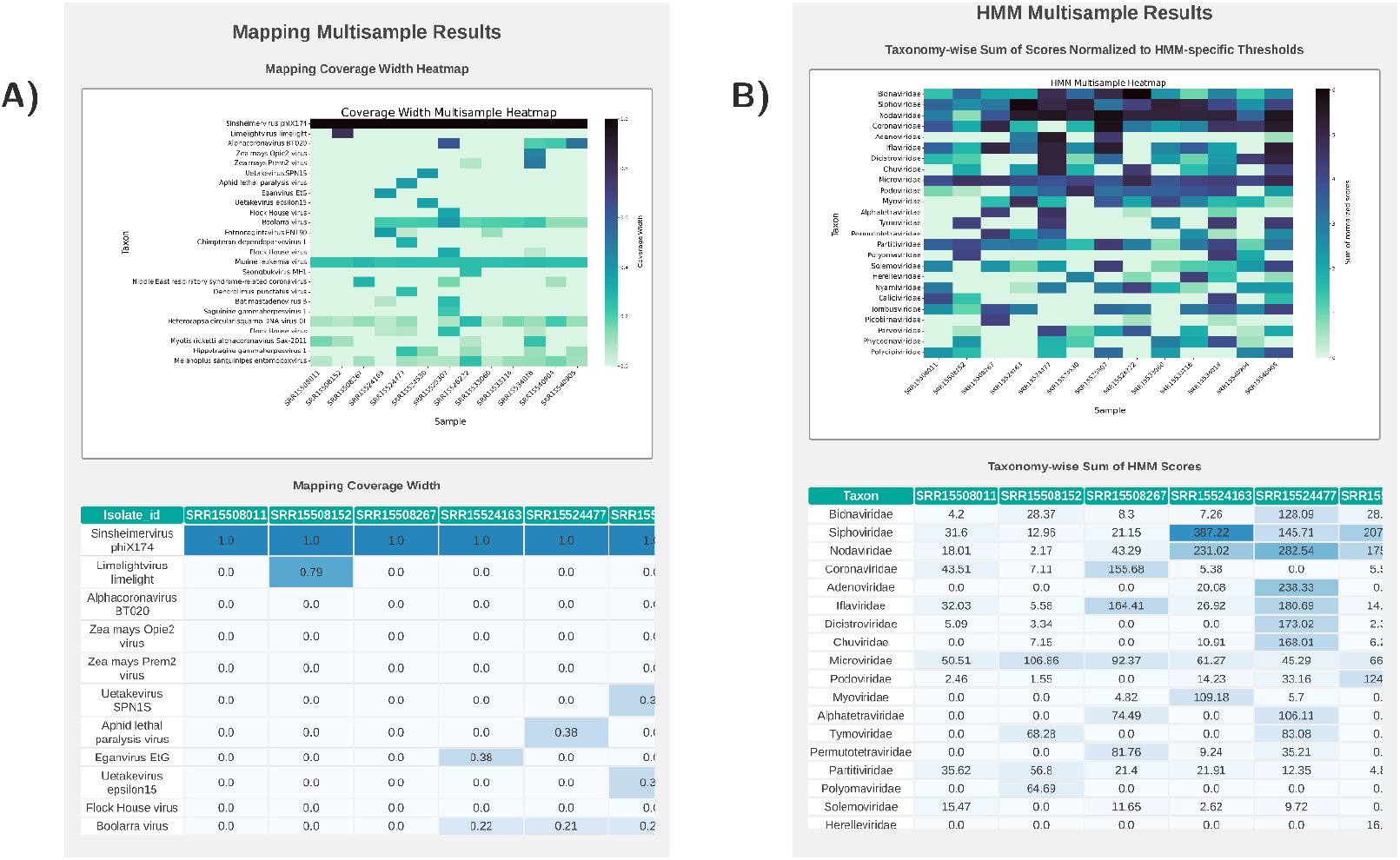
Screenshots of the multisample report of AliMarko. A - a heatmap illustrating coverages across all samples and most represented reference genomes offers a comprehensive view of the findings. Additionally, a scrollable table containing the same information is included. B - The HMM module’s findings are detailed, featuring a heatmap displaying normalized scores and a corresponding results table.

### Applying the pipeline to RNA metagenomic data

We applied AliMarko to the RNA metagenomic data from feces of bats captured in Moscow District of Russia in 2015 (see Methods) [27].

In our analysis of RNA sequencing data from bat fecal samples using AliMarko, we identified the presence of over 25 distinct viruses from animals, plants and bacteria in the dataset. Of these, 7 were mammalian viruses, including representatives from the *Alphacoronavirus, Dependoparvovirus, Betacoronavirus, Mastadenovirus, Sapovirus, Orthopicobirnavirus,* and *Alphapolyomavirus* genera. Notably, more than 2 of the identified viral contigs had lengths approaching those of complete genomes for their respective families. Additionally, 2 contigs of insect viruses had lengths similar to those of the corresponding family genomes (*Iflaviridae* and *Alphatetraviridae*).

In our dataset, we discovered viruses from mammals, insects, and plants. Notably, we found a complete genome of Alphacoronavirus in sample SRR15525307. Furthermore, multiple fragments of *Alphacoronavirus* and *Betacoronavirus* were identified in several samples.

Additionally, a genome of *Caliciviridae* was detected in sample SRR15534018 (Figure 4, A). Using AliMarko, we were able to rapidly determine the presence of Caliciviridae sequences, predict the encoded proteins, and perform phylogenetic analysis of two coding sequences. The phylogenetic analysis revealed that the genome belongs to the Sapovirus clade, although it displays a significant genetic distance from the represented VMR sequences (Figure 4, B). Additionally, the genome’s length and polyA tail, characteristic of this family, suggest that it is likely complete [28]. As a member of the Sapovirus genus, known to cause acute gastroenteritis in humans, the possibility of transmission of viruses from this genus from one host to another is being investigated [29]. Further characterization of the identified Sapovirus contig through a BLASTn search against the nt database revealed only five matches, with the closest homolog being the genome sequence of Bat calicivirus isolate BtCV/OV-157/M.dau/DK/2018 from Denmark (MZ218056.1), sharing 70.6% nucleotide identity with the identified contig [30].

**Figure 4.**
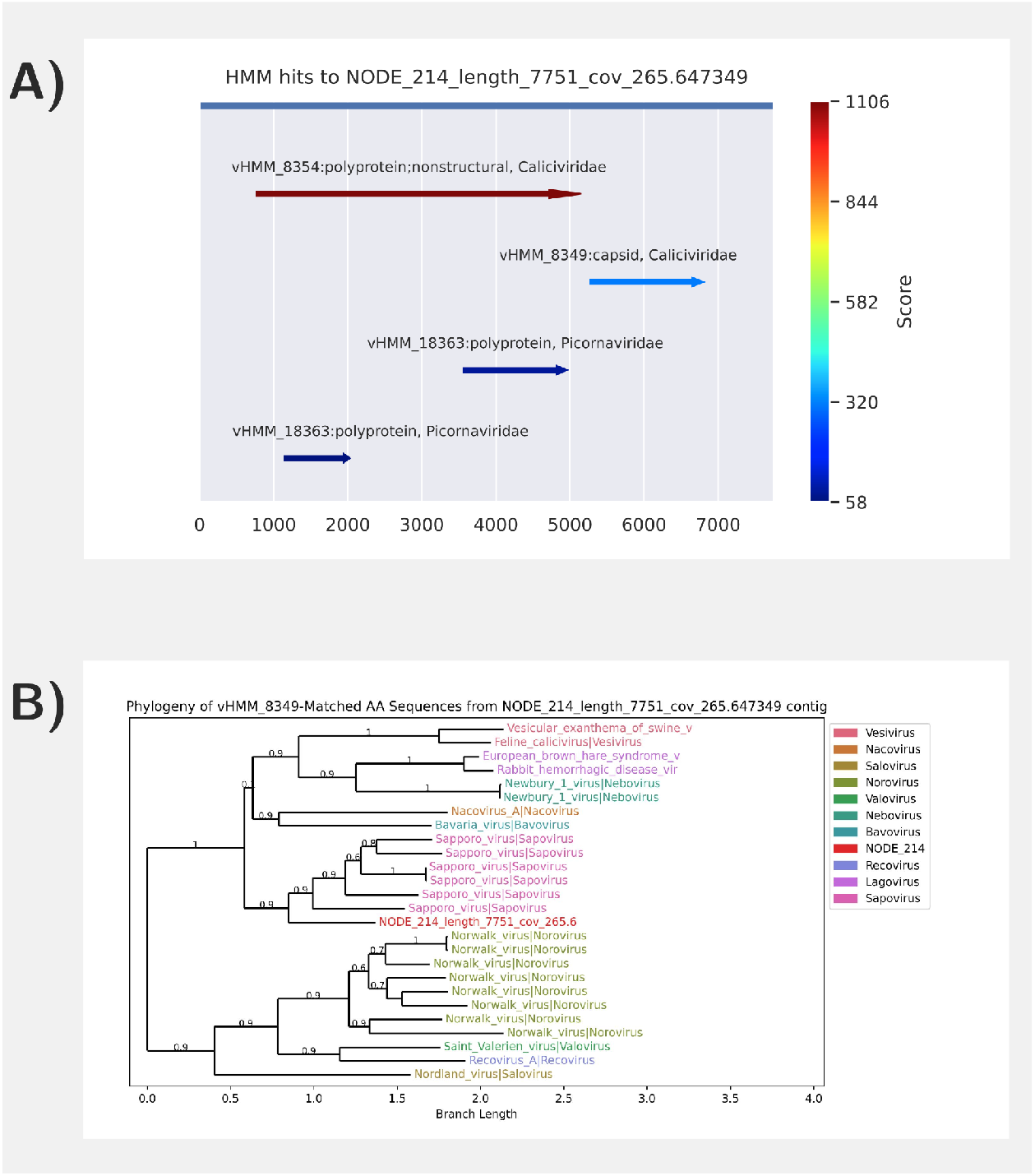
Visualizations from the AliMarko report for SRR15534018 sample. A- visualization of matches of HMM against the Sapovirus contig. The matches are colored by their score. Several models matched the contig. B - phylogenetic tree of contig of presumably Sapovius origin.(see C). Sequences in the tree are colored with their taxonomic group.

Using AliMarko, we identified contigs of the Orthopicobirnavirus genus, which are associated with gastroenteritis in humans. Two contigs were detected, one containing RNA-dependent RNA polymerase sequences (polymerase contig) and the other containing capsid sequences (capsid contig), both of which demonstrated phylogenetic closeness to the Picobirnaviridae family. Notably, phylogenetic analysis with AliMarko revealed that the polymerase contig has high similarity to known sequences, whereas the capsid contig has low similarity. Further analysis using BLAST against the nr database revealed homology with mammalian Picobirnaviridae [31], with the polymerase contig showing 81% nucleotide identity to MK064213.1 [32] and the capsid contig showing 54% identity to WWV86618.1, although with gaps and incomplete coverage [33].

Using a mapping module, we identified sequences in all samples that mapped to the Moloney murine leukemia virus (Retroviridae). The mapping pattern and single-nucleotide substitutions were identical across all samples, suggesting that the source of these sequences is likely a contaminant. Indeed, a previous study (Asplund et al., 2019) has indicated that Nextera kits can serve as a source of these sequences [12].

### Comparison of two methods

In our study, we assessed the diversity of results from two methods. By utilizing our standard databases, we identified unique families. Notably, we processed the results to represent each family once, even if multiple contigs or genome references of a family were involved. Specifically, we focused on families present in both the MINION DB [34, 35] and ICTV VMR databases. Comparing the outcomes revealed that the HMM module with MINION DB detected a greater number of families compared to the alignment module with the ICTV VMR genomes collection (Figure 5). Additionally, a significant number of alignment-identified families were also detected by the HMM module. In general, the HMM module finds two times more families than the alignment module (2.4 for the dataset considered).

**Figure 5.**
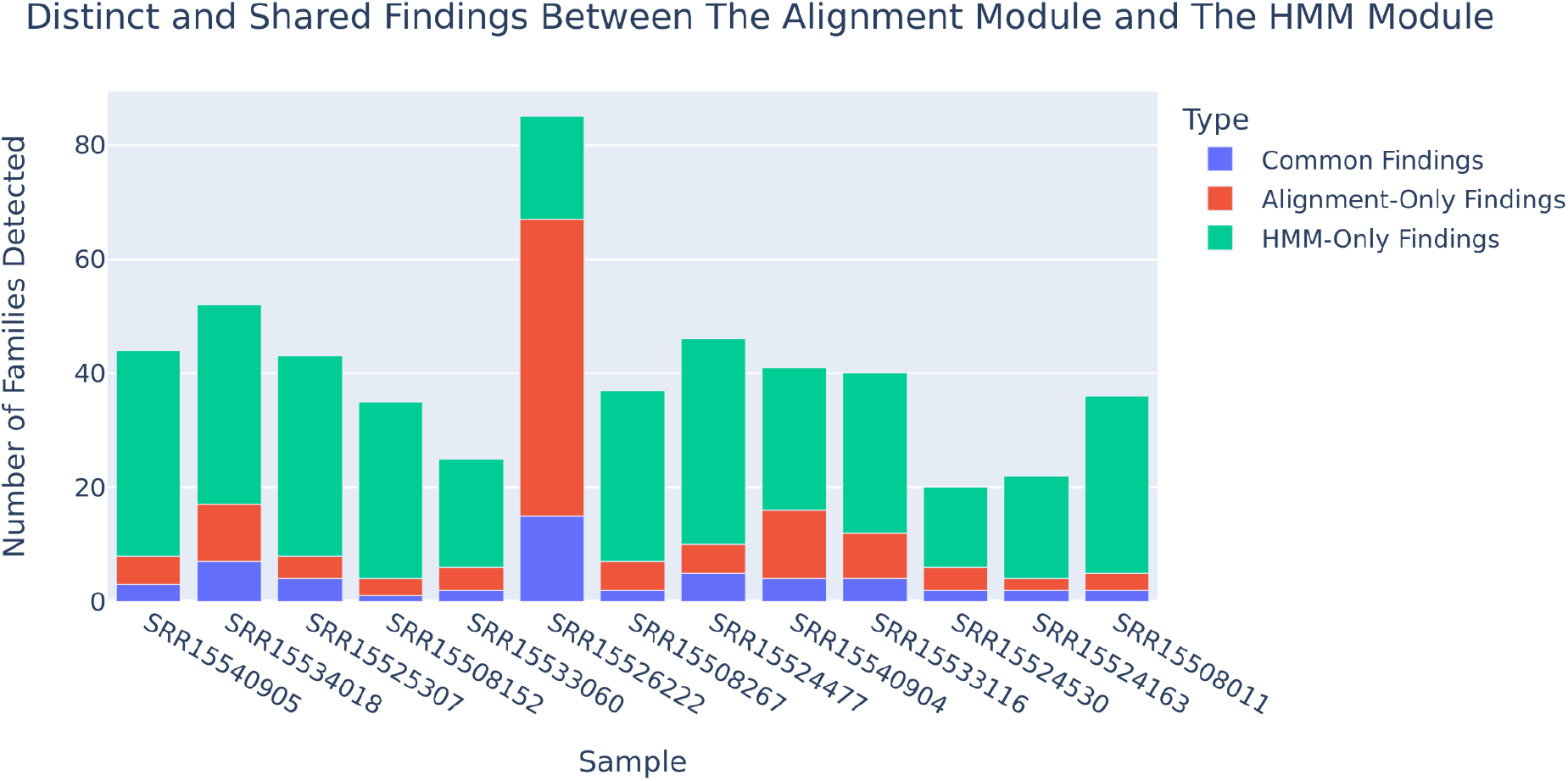
Comparative analysis of Findings by HMM and Alignment modules.

## Discussion

Viruses are a diverse group of organisms, and the absence of universal single-copy genes in viruses presents a significant challenge for identifying viral sequences in metagenomic samples. Additionally, the low abundance of viruses often results in a low concentration of viral nucleic acids in these samples. This makes it difficult for researchers to assemble de novo genomes for most studied viruses, necessitating the use of more sensitive methods [36]. On the other hand, the similarity between sequences of viral and cellular organisms can lead to false positive identifications of viruses. This requires researchers to be cautious when interpreting their results and verifying their validity. Since researchers often need to process large datasets, a pipeline that enables the review of large datasets and performs essential steps to estimate the confidence of individual findings would be valuable.

A core feature of AliMarko is its focus on visualization and interpretable results. The visualizations should enable experts to assess the reliability of findings and distinguish them from potential false positives. The AliMarko pipeline employs two complementary modules: mapping to the reference sequences and de-novo assembly followed by HMM analysis.

The mapping module performs aligning reads to reference sequences from ICTV VMR database and is capable of detecting viral sequences even when the read coverage is too low to assemble contigs.

Evaluating the uniformity of coverage and composition of single nucleotide variants can help researchers estimate the confidence of mapping. If the coverage is uneven, this may suggest to the researcher that the mapping has occurred non-specifically, possibly due to contamination from other organisms. Additionally, identical single nucleotide variant patterns across multiple samples may indicate contamination.

During the analysis of a real dataset, we observed that Murine Leukemia Virus sequences were present in all samples, exhibiting non-uniform coverage across the genome. Each sample had a distinct genomic region covered, with a consistent pattern of nucleotide substitutions. A wide study of linkage between viral sequences and sequencing reagents by Asplund et al. [12] indicated that Murine Leukemia Virus sequences in our dataset are probable contaminants derived from the Nextera kit, which was used to generate our dataset. To avoid false virus identifications, we incorporated the table of potential contaminants provided by Asplund et al. into our pipeline.

In a module complementary to the mapping, the HMM Analysis Module performs read assembly into contigs, amino acid translation and subsequent analysis using HMM and phylogenetic analysis with selected database sequences. This module is effective in identifying viral sequences assembled into contigs of several hundred bases and more. Furthermore, the phylogenetic analysis enables easy interpretation of the result reliability and relationships with reference sequences. In addition to whole-genome phylogenetic analysis, the HMM Analysis Module also enables phylogenetic analysis of individual proteins. This approach allows for consideration of potential differences in mutation accumulation rates between different proteins, providing a more nuanced understanding of the evolutionary relationships between known viral strains and the finding. Future development of this module will focus on integrating automatic topology reliability determination and contig classification to specific clades, enhancing the pipeline’s capabilities in viral sequence analysis.

We applied our pipeline to metagenomic RNA sequencing data from bat fecal samples. Bat metagenomic data are characterized by an extraordinary abundance of viruses, with numerous unknown viruses revealed through sequencing. This is likely due to the unique characteristics of bats, which make them a natural hub for a diverse range of viruses. Our AliMarko-based analysis enabled the detection of over 25 distinct viruses, including seven mammalian viruses from different genera. Notably, we successfully identified a complete genome of a Sapovirus representative. Some members of this genus are known to cause acute gastroenteritis in humans.

Future developments of our pipeline will involve the incorporation of a database of non-viral sequences that are likely to be misidentified as viral (e.g., topoisomerase sequences). This will enable the detection of false mappings to viral sequences and facilitate the rooting of phylogenetic trees. Additionally, Furthermore, we plan to establish a web server hosting AliMarko, providing a user-friendly interface for researchers to access and utilize our pipeline.

We believe that AliMarko will enable researchers to uncover viral sequences in metagenomic datasets, including both viruses with well-understood genomes and those that are novel or unrepresented in existing databases.

## Methods

### Read filtration

Kraken2 is used to deplete reads of cellular organisms in order to lower false-positive viral identifications. We built a custom database that includes the “UniVec_Core”, “archaea”, “bacteria”, “fungi”, “human”, “plant”, and “protozoa” libraries, and it is available as part of AliMarko. Kraken2 is launched with a confidence parameter of 0.7 and the parameter “--unclassified-out” to write unclassified reads to fastq files for further analysis.

Reads deduplication is performed with fastp [37]. Reads with quality scores less than 15 are excluded using samtools view.

### Read mapping and analysis

Read mapping is performed with BWA-MEM [38]. Mappings with a quality score of less than 20 and match percentage less than 63% are eliminated. Coverage width and mean mapping quality of mapping for each reference genome are calculated with samtools.

Mapping visualization is performed with BamSnap [39]. We modified the source code of BamSnap to allow it to work with full visualization of short sequences. Modified version is available at: https://github.com/NJJeus/bamsnap.

### HMM analysis

De-novo assembly is performed using SPAdes [40] with the -meta option. HMM analysis is conducted using the pyHMMer library [41] and Viral Minion DB HMM [34, 35] on six-frame translated nucleotide sequences.

For each HMM, a threshold score was determined as follows. All non-viral proteins (negative set) were selected from the Swiss-Prot database [42]. The hmmscan tool was then run on this set, and the highest score obtained was designated as the threshold for further use. When performing analysis, only HMM matches with a score greater than the threshold are further considered. Regions of the contig that match the HMM profile are visualized using custom Python code.

### Phylogenetic analysis

The phylogenetic analysis is conducted as follows: a region identified using HMM is extracted from the contig. Additionally, sequences from reference genomes are analyzed using the same HMM, and the identified regions are extracted. All these extracted regions are then added to a FASTA file. Multiple sequence alignment is performed using MAFFT [43]. A phylogenetic tree is constructed with FastTree [44], which infers approximately maximum-likelihood phylogenetic trees. FastTree provides local support values based on the Shimodaira-Hasegawa test to estimate the reliability of each split in the tree, which are depicted in a tree visualization.

### Task management

The pipeline is written in Snakemake, a modern workflow management system that allows for simultaneous analysis of multiple samples, manages the use of multiple threads, and facilitates the installation of all required libraries.

### Materials

We tested our pipeline on total NGS RNA sequencing of fecal samples from 26 bats captured in the Zvenigorodsky District of the Moscow Region [27]. RNA was extracted from the bat fecal samples using the QIAamp Viral RNA Mini Kit (Qiagen, Germany). The extracted RNA was used for reverse transcription with Reverta-L reagents (AmpliSens, Russia), and second-strand cDNA was synthesized using the NEBNext Ultra II Non-Directional RNA Second Strand Module (New England Biolabs, E6111L). Sequencing was performed on the Illumina MiSeq system to generate 250-bp paired-end reads. The data are available in SRA database by accessions SRR15540905, SRR15540904, SRR15534018, SRR15533116, SRR15533060, SRR15526222, SRR15525307, SRR15524477, SRR15524163, SRR15524530, SRR15508152, SRR15508267, SRR15508011.

To demonstrate the functionality of our computational pipeline, we created a small simulated dataset. This dataset comprised short reads emulating HS25 sequencing, generated using the art_illumina simulator [45]. The reference genomes used for simulation included a selection of bacterial species, the human genome, and the genomes of two viruses: Xanthomonas phage phiXv2 and Miniopterus bat coronavirus 1 (Table S1). Additionally, we incorporated sequences identified as belonging to Murine leukemia virus, which were derived from bat sequencing data.

## Supplementary

**Table S1**: Genome references for simulated sample

**HTML S1.** An example of AliMarko sample report on a simulated sample

## Supporting information

Table S1: Genome references for simulated sample

HTML S1. An example of AliMarko sample report on a simulated sample

