## Supplementary material for "AliMarko: A Novel Tool for Eukaryotic Virus Identification Using Expert-Guided Approach": HTML S1. An example of AliMarko sample report on a simulated sample: HTML S1. Simulated Sample.html

 

Simulated Sample


### Sample: simulated sample

#### Mapping Summary

| Species | Host source | Coverage width | Mean depth | Genus | Family | Realm |
| --- | --- | --- | --- | --- | --- | --- |
| Miniopterus bat coronavirus 1 | vertebrates | 1.0 | 19.91 | Alphacoronavirus | Coronaviridae | Riboviria |
| Lophivirus Xv2 | bacteria | 0.99 | 19.56 | Lophivirus | Inoviridae | Monodnaviria |
| Murine leukemia virus | vertebrates | 0.24 | 6.57 | Gammaretrovirus | Retroviridae | Riboviria |

#### HMM Hit Summary

| HMM | Taxon | Name | Putative Protein | Score |
| --- | --- | --- | --- | --- |
| vHMM\_8398 | Coronaviridae | NODE\_1\_length\_28274\_cov\_12.656579 | spike;glycoprotein | 1568 |
| vHMM\_8393 | Coronaviridae | NODE\_1\_length\_28274\_cov\_12.656579 | replicase;polyprotein;1ab | 6126 |
| vHMM\_8393 | Coronaviridae | NODE\_1\_length\_28274\_cov\_12.656579 | replicase;polyprotein;1ab | 4939 |

---

#### Mapping Details

#### Host: bacteria

##### Lophivirus Xv2

| Fragment | Len | Coverage width | Nucleotide similarity | Mean Depth | MeanMAPQ | SNP Count |
| --- | --- | --- | --- | --- | --- | --- |
| MH206183 | 6564 | 0.9906 | 1.0 | 19.5612 | 0.593 | 0.0 |

#### Host: vertebrates

##### Miniopterus bat coronavirus 1

| Fragment | Len | Coverage width | Nucleotide similarity | Mean Depth | MeanMAPQ | SNP Count |
| --- | --- | --- | --- | --- | --- | --- |
| EU420138 | 28326 | 0.9982 | 1.0 | 19.911 | 0.6 | 0.0 |

##### Murine leukemia virus Sequences of this virus have association with Nextera.

| Fragment | Len | Coverage width | Nucleotide similarity | Mean Depth | MeanMAPQ | SNP Count |
| --- | --- | --- | --- | --- | --- | --- |
| AF033811 | 8332 | 0.2439 | 1.0 | 6.5653 | 0.586 | 0.0 |


---

#### HMM Module Results by Contig

#### Contigs

##### NODE\_1\_length\_28274\_cov\_12.656579:Coronaviridae

| HMM | Taxon | Name | Putative Protein | Score |
| --- | --- | --- | --- | --- |
| vHMM\_8398 | Coronaviridae | NODE\_1\_length\_28274\_cov\_12.656579 | spike;glycoprotein | 1568 |
| vHMM\_8393 | Coronaviridae | NODE\_1\_length\_28274\_cov\_12.656579 | replicase;polyprotein;1ab | 6126 |
| vHMM\_8393 | Coronaviridae | NODE\_1\_length\_28274\_cov\_12.656579 | replicase;polyprotein;1ab | 4939 |

##### Phylogenetic Tree of vHMM\_8393-Matched Amino Acid Sequences

##### Putative protein: replicase;polyprotein;1ab

##### Phylogenetic Tree of vHMM\_8398-Matched Amino Acid Sequences

##### Putative protein: spike;glycoprotein

##### Phylogenetic Tree of vHMM\_8393-Matched Amino Acid Sequences

##### Putative protein: replicase;polyprotein;1ab
